# Numerosity adaptation suppresses early visual responses

**DOI:** 10.1101/2024.08.29.610311

**Authors:** Liangyou Zhang, Evi Hendrikx, Yizhen Wang, Surya Gayet, Serge O. Dumoulin, Ben M. Harvey

## Abstract

Humans and many animals rapidly and accurately perceive numerosity, the number of objects, in a visual image. The numerosity of recently viewed images influences our perception of the current image’s numerosity: numerosity adaptation. How does numerosity adaptation affect responses to numerosity in the brain? Recent studies show both early visual responses that monotonically increase with numerosity, and later numerosity-tuned responses that peak at different (preferred) numerosities in different neural populations. We have recently shown that numerosity adaptation affects the preferred numerosity of numerosity-tuned neural populations. We have also shown that early visual monotonic responses reflect image contrast, which follows numerosity closely. Here we ask how monotonic responses in the early visual cortex are affected by adaptation to different numerosities, using ultra-high field (7T) fMRI and neural model-based analyses. FMRI response amplitudes increased monotonically with numerosity throughout the early visual field maps (V1-V3, hV4, LO1-LO2 & V3A/B). This increase in response amplitudes becomes less steep after adaptation to higher numerosities, with this effect becoming stronger through the early visual hierarchy. This suppression of responses to numerosity is consistent with perceptual effects where adaptation to high numerosities reduces the perceived numerosity. These results imply that numerosity adaptation effects in later numerosity-tuned neural populations may originate in early visual areas that respond to image contrast in the adapting image.

## Introduction

Numerical cognition leverages aspects of perception, attention and working memory to construct a quantitative understanding of the world that eventually allows advanced abilities like mathematics. Numerical cognition’s simplest stages require only an ability to estimate and perceive object number, or numerosity, often called the ‘number sense’. This simple numerosity perception is found in many animals, including humans (Burr & Ross, 2008; Jevons, 1871) and non-human primates (Nieder et al., 2002), but also birds (Scarf et al., 2011), amphibians (Krusche et al., 2010), fish (Miletto Petrazzini et al., 2016), and insects (Howard et al., 2018). Numerosity perception may provide a selective advantage in any animal by helping to forage for food, like finding the plant with the most fruits (Nieder, 2021, 2022). Some have proposed that numerosity perception reflects non-numerical image features that are often correlated with numerosity, like density (Durgin, 2008) or contrast energy at high spatial frequencies (Dakin et al., 2011). However, extensive recent evidence demonstrates that humans perceive numerosity itself more quickly and accurately than these non-numerical features (Cicchini et al., 2016; DeWind et al., 2015; Testolin et al., 2020).

Which neural responses underlie numerosity perception? Two broad classes of responses have been described: numerosity-tuned responses and monotonically changing responses. In numerosity-tuned neural populations, the response peaks at a specific (preferred) numerosity and gradually decreases with distance from this numerosity (Nieder et al., 2002) (Nieder & Miller, 2003). Numerosity-tuned responses have been found in single neurons in monkeys (Nieder et al., 2002; Nieder & Miller, 2003), humans (Kutter et al., 2018), crows (Ditz & Nieder, 2016) and chickens (Kobylkov et al., 2022). We have also revealed the numerosity tuning of neural populations throughout the human brain by combining ultra-high field (7T) functional magnetic resonance imaging (fMRI) and neural model-based analyses (Cai et al., 2021; Harvey et al., 2013; Harvey & Dumoulin, 2017). Numerosity-tuned responses are located in the association cortices of humans (Harvey & Dumoulin, 2017) and monkeys (Nieder & Miller, 2004). Their responses closely predict numerosity perception across trials (Nieder & Miller, 2003; Tudusciuc & Nieder, 2007) and across individuals (Kersey & Cantlon, 2017; Lasne et al., 2019; Piazza et al., 2004), positioning them as an important basis of numerosity perception (Tsouli et al., 2022).

The second class of neural populations increase their response amplitude monotonically as numerosity increases. Such monotonic responses, sometimes called summation coding (Roggeman et al., 2007; Zorzi & Testolin, 2018), have long been predicted as an intermediate stage in computational models for the derivation of downstream numerosity-tuned responses (Dehaene & Changeux, 1993; Kim et al., 2021; Stoianov & Zorzi, 2012; Verguts & Fias, 2004; Zorzi & Testolin, 2018). Monotonically responding populations have been described using EEG and fMRI (DeWind et al., 2019; Park et al., 2015; Paul et al., 2022). These responses were found in early visual cortex (including V1 (Paul et al., 2022), V2, and V3 (Fornaciai & Park, 2018a; Paul et al., 2022)) with very short latencies (Park et al., 2015), indicating that these monotonic responses are implicated in the earlier stages of feedforward visual processing. Surprisingly for early visual responses, these responses to numerosity are not strongly affected by non-numerical features like item size and spacing (DeWind et al., 2019; Park et al., 2015; Paul et al., 2022). How can such early responses already encompass numerosity information?

These early visual responses may be explained by close relationships between numerosity and aggregate Fourier power in the stimuli used in most experiments (Paul et al., 2022). This aggregate Fourier power follows numerosity closely, with little effect of item size or spacing. Indeed, monotonically increasing responses in early visual areas follow the logarithm of aggregate Fourier power more closely than the logarithm of numerosity (Paul et al., 2022). As such, the established response properties of the early visual cortex give a population-level monotonic response to Fourier power from which numerosity can be straightforwardly computed. This allows for numerosity-tuned populations to emerge in lateral occipital cortex (Paul et al., 2022) and propagate throughout the association cortices. In sum, while numerosity estimation can be described as a relatively simple perceptual ability beginning at the earliest stages of vision, the resulting numerosity representation may be used for higher-order cognitive processes throughout the brain (Harvey & Dumoulin, 2017; Paul et al., 2022).

Like many visual features, numerosity perception is affected by adaptation (Burr & Ross, 2008), where perceived numerosity is repelled from previously-presented numerosities. For example, after repeatedly viewing a high numerosity, lower numerosities are underestimated. How does numerosity adaptation affect neural responses to numerosity? First, in fMRI repetition suppression paradigms, repeated presentation of a single numerosity suppresses parietal neural responses to similar numerosities more than responses to more different numerosities (Piazza et al., 2004). This is seen as evidence for numerosity-tuned responses in human parietal cortex and suggests that adaptation strongly affects numerosity-tuned responses. Similarly, numerosity adaptation strongly reduces the ability to distinguish between the patterns of activity evoked by different numerosities in parietal cortex using multivariate classification methods, suggesting adaptation suppresses or changes the patterns of response to specific numerosities (Castaldi et al., 2016; Eger et al., 2009). Finally, we have shown that numerosity tuning in neural populations with numerosity preferences near the adaptor is repelled from the adapted numerosity while that in neural populations with numerosity preferences further from the adaptor is attracted toward the adapted numerosity (Tsouli et al., 2021). This mixed change in numerosity preferences occurs in all numerosity-tuned responses throughout the association cortices and suggests some form of normalization across the whole set of numerosity-tuned responses.

However, it remains unclear whether early visual monotonic responses to numerosity are affected by adaptation and may therefore contribute to later adaptation effects on numerosity-tuned responses. Here, we therefore analyzed these early visual monotonic responses in an ultra-high field (7T) fMRI data set where we have previously shown numerosity adaptation effects on numerosity-tuned responses (Tsouli et al., 2021). During fMRI scanning, participants viewed the same changing sequence of numerosities (to map numerosity preferences) under conditions of low numerosity adaptation, high numerosity adaptation and changing numerosity adaptation. In the current study, we compared the amplitudes of responses to these conditions in the early visual cortex. Higher numerosities produce a stronger neural response in early visual cortex than lower numerosities. We therefore hypothesized that adaptation to a higher numerosity would more strongly suppress the monotonic response to subsequently viewed displays, by more strongly reducing the sensitivity of the responsive neural populations.

## Results

### Monotonic responses to numerosity displays in the early visual cortex

During fMRI scanning, participants viewed sequences of progressively increasing and decreasing numerosities (from one to seven and back) to quantify response amplitudes to different numerosities. These progressively changing numerosity displays were presented in three different adaption conditions (Fig. 1): 1) Preceded by displays containing one item (low adaptor condition); 2) Preceded by displays containing twenty items (high adaptor condition); 3) Preceded by displays of the same changing numerosity (changing adaptor condition). In the changing adaptor condition, the changing numerosity was therefore presented twice as frequently as other conditions and so will produce larger monotonic responses.

**Figure 1:**
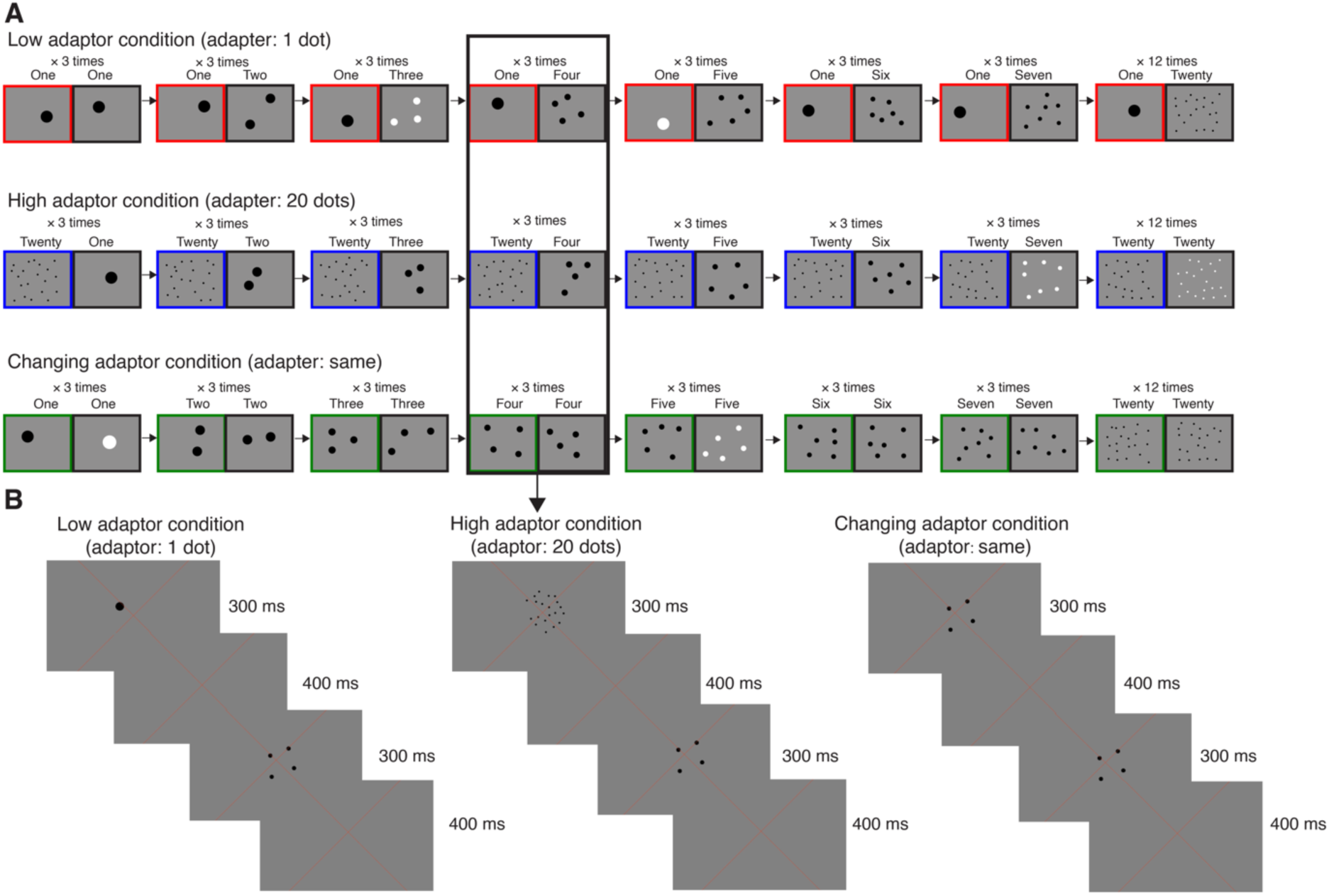
Numerosity stimuli. (A) Schematic description of the numerosity response mapping stimuli shown in the ascending progression of one stimulus cycle. Each fMRI time frame (TR) contained an adaptor numerosity (left, colored border), which differed between conditions, followed by a changing numerosity (right, black border). In all three conditions the changing numerosities increases from 1 through 7, followed by a baseline of 20 dots. In the low and high adaptor conditions, the adaptor was constant at numerosities of 1 and 20 respectively. In the changing adaptor condition, the changing numerosities were also shown as the adaptor. In all conditions, the same pair of adaptor and changing numerosities was repeated three times (across three TRs) to ensure strong fMRI responses. (B) Example displays presented in a single TR in each condition.

We used a changing adaptor condition (Fig. 1), without adaptation to a fixed numerosity, to identify responses to changes in numerosity as we have used this stimulus design in previous studies (Harvey et al., 2013; Harvey & Dumoulin, 2017; Paul et al., 2022; Tsouli et al., 2021) and it maximizes the neural response amplitude and goodness of fit of our models. We explained responses in all conditions using a monotonic response model where the amplitudes of the neural response underlying the fMRI response was proportional to the logarithm of the aggregate Fourier power of the changing numerosity display shown during each fMRI time frame. As previously shown (Paul et al., 2022), many recording sites showed such responses in the representation of the central visual field (where the numerosity mapping stimulus was displayed) in visual field maps V1-V3, hV4, V3A/B, LO1 and LO2 (Fig. 2). We selected recording sites in each visual field map where preferred visual position eccentricity was below 1°, where a monotonic response increasing with aggregate Fourier power explained at least 10% of response variance, and where this monotonic response model explained more variance than a numerosity-tuned model.

**Figure 2:**
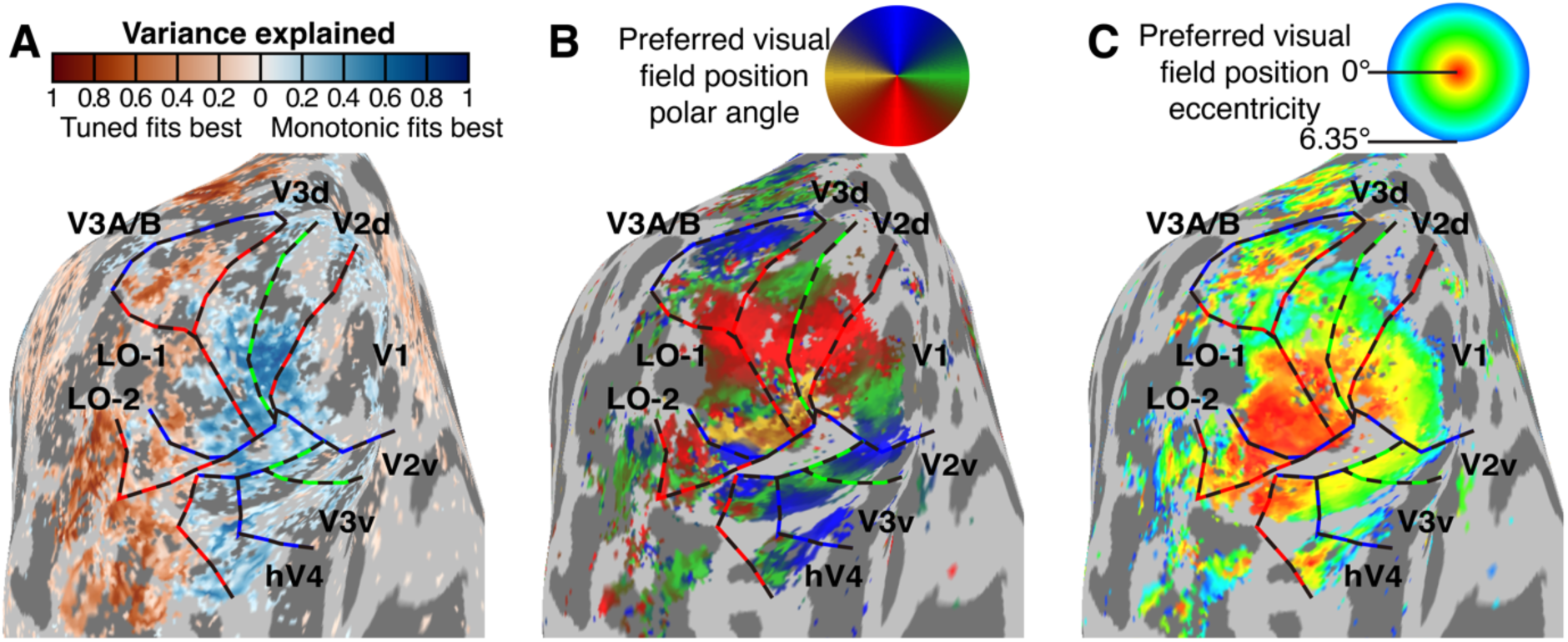
Locations of monotonic responses to numerosity. (**A**) Blue recording sites show responses that monotonically increased in proportion to the logarithm of aggregate Fourier power, while red recording sites show numerosity-tuned responses. Here, the best-fitting response model explained at least 0.1 (cross-validated R^2^) of response variance. (**B**) The preferred visual field position polar angle of each recording site (obtained from visual field mapping data) let us localize visual field map borders at reversals in polar angle progressions. Dashed lines show visual field map borders at the upper vertical meridian (blue) lower vertical meridian (red) and horizontal meridian (green). (**C**) Each recording site’s preferred visual field position eccentricity. We used this to localize sites with a preferred eccentricity below 1°, whose population receptive fields included the numerosity mapping stimulus area.

### Changes in early visual monotonic responses during numerosity adaptation

We first asked whether the change in response amplitudes over the course of a scan in the recording sites differed between the adaptor conditions. The fMRI responses of recording sites in the early visual cortex increased following the aggregate Fourier power (and so the numerosity) of the presented displays in all adaptation conditions (Fig. 3A). However, the monotonic change in response amplitudes was greater in the low than in the high adaptor condition. Note that this change in response amplitude was greater still in the changing adaptor condition. A likely explanation for this is that the adaptor in the changing adaptor condition contains the same changes in numerosity as the changing numerosity stimulus does. This effectively doubles the changes in numerosity between fMRI time frames. For example, the difference between the total numerosity in the first and second displays in Fig. 1 is two ((2+2)-(1+1)) in the changing adaptor condition but only one in the high adaptor condition (20+2)-(20+1) and low adaptor (1+2)-(1+1) conditions.

**Figure 3.**
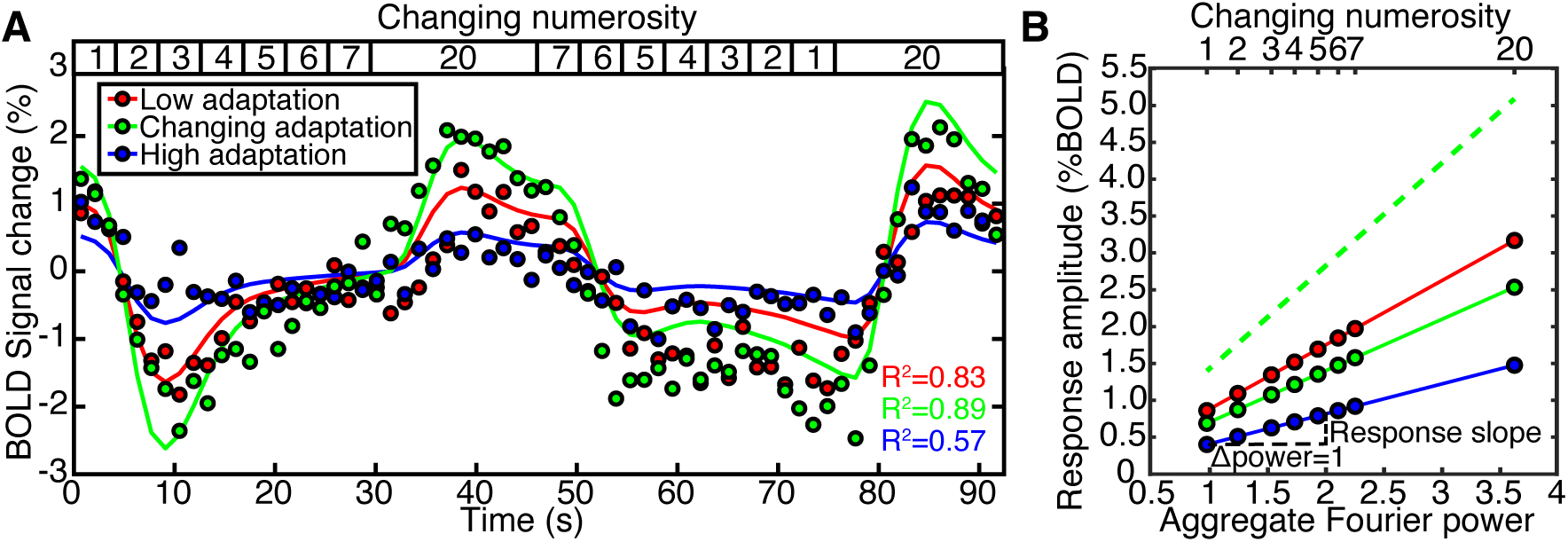
The response of an example recording site (voxel) in V1 to a numerosity mapping stimulus differs with numerosity adaptation. (**A**) As the stimulus’ changing numerosity progressively increased and decreased (top inset), the fMRI BOLD response in all adaptation conditions (colored dots) increased and decreased, after a hemodynamic delay. The responses in all conditions were closely fit by the predictions of the monotonic responses to the aggregate Fourier power of the stimulus (colored lines), scaled with different amplitudes. The range of response amplitudes was greater in the low adaptor condition than the high adaptor condition. The range of response amplitudes in the changing adaptor condition was greater still, because both the adaptor numerosity and the changing numerosity changed in the same way so the changing numerosity is presented twice as frequently. The variance explained (R^2^) followed this range of response amplitudes, as a lower amplitude decreases the signal-to-noise ratio of the response. (**B**) We explained these responses using neural response models in which neural responses monotonically increase proportionally to the aggregate Fourier power of the displays, which follows numerosity closely (Paul et al., 2022). We fit the slope of this proportionality (i.e. the increase in amplitude of the neural response when aggregate Fourier power increases by one, Δpower=1) using a general linear model.

In our response model, the response amplitude of each recording site in each condition was captured by a slope (beta) parameter. This quantified how much the amplitude of the neural response underlying the fMRI signal increased when the logarithm of the aggregate Fourier power of the stimuli increased by one (Fig 3B). Our response model included the effect of the adaptor stimuli. This was constant in the low and high adaptor condition, so could not explain any response variance, but changed with the changing numerosity in the changing adaptor condition. After including responses to these changes, the amplitude slope in the changing adaptor condition fell between those of the low and high adaptor conditions. This slope was greatest in the low adaptor condition, intermediate in the changing adaptation condition, and smallest in the high adaptor condition. In a response model of the changing adaptor condition that ignored responses to the adaptor (green dashed line), the slope doubled compared to the model that considered responses to the adaptor (green solid line).

To summarize the response change in each visual field map, we first calculated the response slope in each adaptation condition for recording sites throughout the early visual cortex (from V1 to LO2 and V3A/B) (Fig 4A-C). Notably, some later visual field maps (LO1, LO2, V3A/B) include recording sites with both monotonic and tuned responses: our analyses included only recording sites whose responses are better explained (better fit under cross-validation) by our monotonic response model in the changing adaptor condition. For each visual field map, we took the average slope in each condition across these recording sites in every hemisphere and used these hemisphere averages for statistical comparisons. The slope was significantly above zero in all adaptation conditions in all visual field maps, so all conditions yielded monotonically increasing response to aggregate Fourier power (and so to numerosity) in all visual field maps (Fig. 4D). In each visual field map, this slope was significantly greater in the low adaptor condition than the changing adaptor condition, which in turn had a greater slope than the high adaptor condition (Fig. 4D). This reduction in slopes from lower to higher adaptor conditions reflects neural adaptation of monotonic responses to increasing numerosities and is also consistent with perceptual numerosity adaptation effects, where adaptation to higher numerosities decreases the perceived numerosity of lower numerosity stimuli.

**Figure 4:**
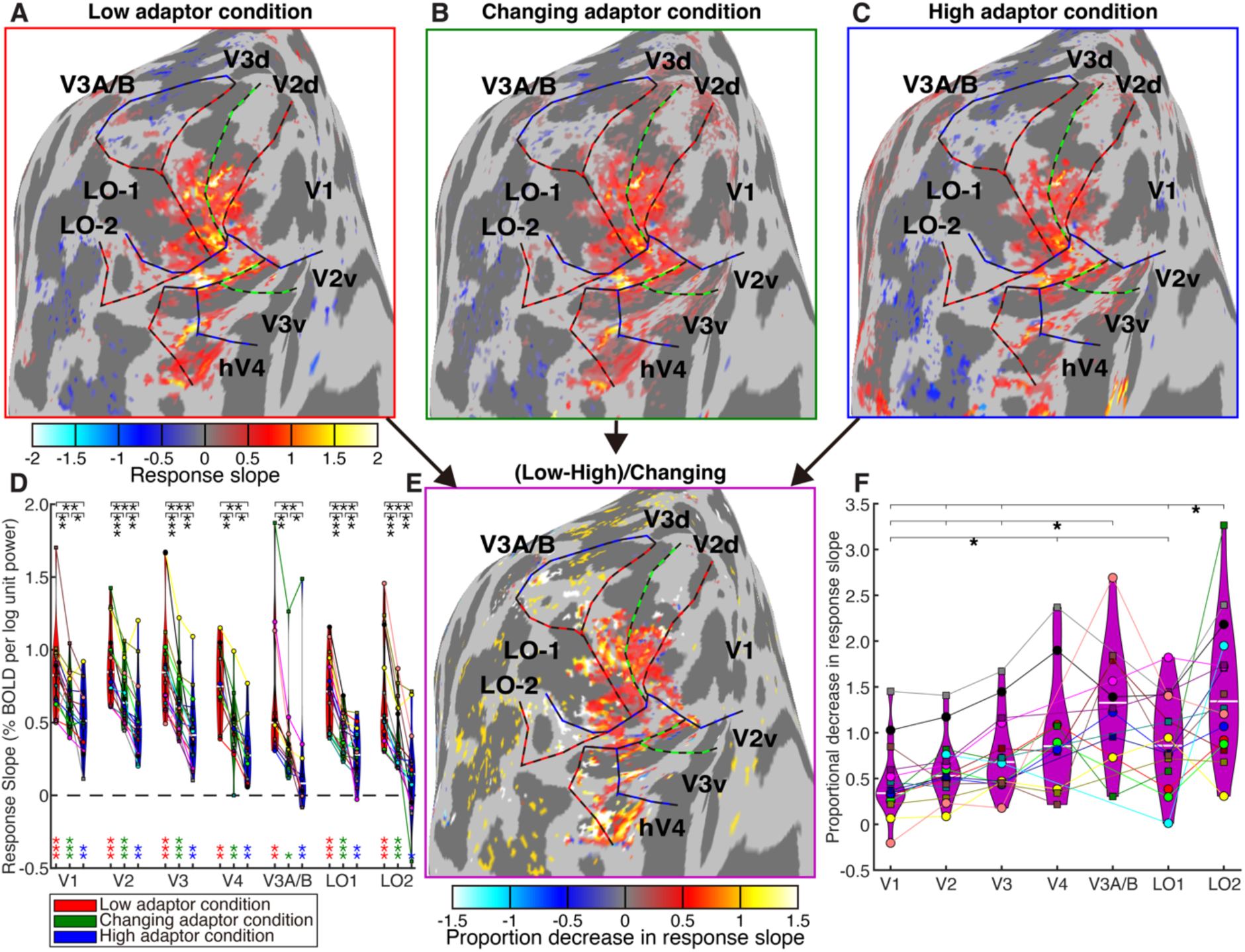
Neural adaptation of monotonic responses increased through the visual hierarchy. (**A-C**) The fMRI BOLD responses increased monotonically with aggregate Fourier power in recording sites throughout the central visual field representations of the early visual field maps, in the low (A), changing (B) and high (C) adaptor conditions. (**D**) In the average across the recording sites in each visual field map of each hemisphere, the slope of the monotonic response increase with the logarithm of aggregate Fourier power was significantly positive in all conditions and all visual field maps (colored stars). This slope was greatest in the low adaptor condition, intermediate in the changing adaptor condition, and lowest in the high adaptor condition (black stars show comparisons between conditions in each visual field map) \**p*<0.05, \*\**p*<0.01, \*\*\**p*<0.001. Colored markers (linked with colored lines) show the mean in the visual field map example in each hemisphere and condition. (**E**) To compare this reduction in monotonic response amplitude between visual field maps, we calculated the change in response slope between adaptation conditions (here: low minus high) in each recording site, as a proportion of the slope in the changing adaptor condition. (**F**) This proportional decrease in response amplitude from low to high adaptor conditions (i.e., the neural adaptation effect strength) became greater through the visual processing hierarchy. Visual field maps marked with brackets to the right of the stars showed significantly stronger proportional decreases than those with brackets to the left of the starts.

### Progressive increases in adaptation effects through the early visual hierarchy

We then compared the strength of this neural adaptation effect between visual field maps. We first computed a measure of relative slope change; specifically, we subtracted the slope in a high adaptor condition from the slope in a low adaptor condition, and divide by the slope of the changing adaptor condition (Fig. 4E). This metric reflects how much shallower the slope (of responses to increasing numerosities) becomes when preceded by a high compared to a low adaptor, quantifying the relative strength of the neural adaptation effect in each visual field map. An ANOVA (with visual field map as a fixed factor and participant as a random factor) found significant differences between maps in this proportional reduction in response slope (*F*(6,85)=11.4, *p*=2.5×10^-9^). Post-hoc multiple comparisons revealed a progressive increase in the strength of the neural adaptation effect from earlier to later visual field maps (Fig. 4F). Therefore, the effect of numerosity adaptation on neural response amplitudes increased through the early visual hierarchy.

## Discussion

In the current study, we asked whether numerosity adaptation affects the responses of the early visual cortex. First, we found that the monotonically increasing neural response to numerosity occurred regardless of numerosity adaptation. These effects began by V1 and continued through the early hierarchy to V2, V3, hV4, V3A/B, and LO1-LO2. Second, in all these visual field maps, the amplitude of this monotonic increase (slope) was reduced when the adapting numerosity was higher. This is consistent with the perceptual effect where perceived numerosity is reduced during high numerosity adaptation. Third, the proportion by which the response slope was reduced during higher compared to lower numerosity adaptation (i.e., the magnitude of the adaptation effect) increased hierarchically from V1 onward.

In this study, we focus on the early visual neural response that monotonically increases with numerosity (DeWind et al., 2019; Park et al., 2015; Paul et al., 2022). We have explained these findings by the close relationship between numerosity and contrast energy in the spatial frequency domain (Paul et al., 2022). At a fixed contrast, this aggregate Fourier power follows numerosity closely but nonlinearly, with little effect of size or spacing, and predicts population responses in V1 and computational models (Kim et al., 2021; Stoianov & Zorzi, 2012) more closely than numerosity does (Paul et al., 2022). This provides a signal from which numerosity itself may be straightforwardly derived. Indeed, responses in the numerosity-tuned populations of the association cortices are more closely predicted by numerosity than aggregate Fourier power (Paul et al., 2022). Therefore, we describe the monotonic responses in the current study as responses to contrast and the tuned responses as responses to numerosity. However, we found a very similar pattern of results if we model the early visual responses as functions of the log(numerosity), rather than as functions of aggregate Fourier power, of our displays.

Adaptation effects on numerosity perception (Burr & Ross, 2008) have always been assumed to reflect changes in the responses of numerosity-tuned neurons. This is a reasonable assumption for several reasons. First, when adaptation of numerosity perception was first described, numerosity-tuned neurons had recently been found in macaque parietal and frontal cortices (Nieder et al., 2002; Nieder & Miller, 2004), and tuned effects of repetition suppression were found in human parietal cortex (Piazza et al., 2004). Early visual monotonic responses to numerosity were only described years later (DeWind et al., 2019; Park et al., 2015). Second, adaptation effects are often found for image features with tuned neural representations, like orientation (Dragoi et al., 2000) and motion direction (Mather, 1980). Third, adaptation to a low numerosity has been shown to increase perceived numerosity (Aulet & Lourenco, 2023; Burr & Ross, 2008), as well as adaptation to high numerosity decreasing perceived numerosity. The bidirectionality of this repulsive effect seems likely to reflect effects on numerosity-tuned neural populations with different numerosity preferences. Specifically, adaptation to a numerosity below the numerosity preference of a numerosity-tuned neuron should suppress responses to lower numerosities more than responses to higher numerosities. This should thereby increase the numerosity yielding the largest response (the numerosity preference) (Tsouli et al., 2022). Accordingly, we have recently shown (using the present data set) that tuned neural numerosity preferences are affected by adaptation (Tsouli et al., 2021).

However, converging evidence also suggests that the neural effects of numerosity adaptation begin at early visual processing stages, with spatially specific responses to image contrast. First, perceptual numerosity adaptation is highly spatially specific (limited to the adapted location) (Burr & Ross, 2008), while numerosity-tuned neurons have large spatial receptive fields and their response to numerosity does not depend on the stimulus falling within that receptive field (Harvey et al., 2015; Harvey & Dumoulin, 2017; Paul et al., 2022; Viswanathan & Nieder, 2020). Second, perceptual numerosity adaptation is weaker when the adaptor and test displays differ in color (Grasso et al., 2022) or other low-level visual features (Caponi et al., 2024). Different low-level features activate distinct neural populations in early visual processing, but similar numerosity-tuned responses are found regardless of item color (Cai et al., 2022). Third, compelling recent results (Bonn & Odic, 2023) show that perceived numerosity is affected by adaptation to gratings with no numerosity but a spatial frequency matching that of the numerosity display. Fourth, recent results show that the strength of the numerosity adaptation effect is greater when the positions of the individual items in the adaptor and test displays overlap (Yousif et al., 2023). Fifth, the increase in perceived numerosity after low numerosity adaptation is far weaker than the decrease after high numerosity adaptation (Aulet & Lourenco, 2023; Yousif et al., 2023). The asymmetry of this bidirectional effect may reflect an additional effect of adaptation at the monotonic response stage for high numerosity displays. Finally, the numerosity adaptation effect becomes weaker as contrast decreases (Burr & Ross, 2008), though it remains clear even at low contrasts. Together with the present results, these results suggest that perceptual numerosity adaptation at least partly originates in early visual processing stages with spatially specific responses to contrast.

Importantly, none of these findings show that perceptual numerosity adaptation arises only through early visual contrast adaptation and indeed several results speak against this interpretation. First, we found that effects on monotonic responses become progressively stronger through the early visual hierarchy. Second, recent results (Kido et al., 2024) show that responses to numerosity in more anterior areas of the association cortices depend progressively more on the context of recently-presented numerosities. Third, the effects on monotonic responses that we see are only correlated with effects on tuned responses in the most posterior numerosity map. All of these results suggest progressively increasing neural adaptation effects throughout the numerosity processing hierarchy, not effects at an early stage alone. Furthermore, adaptation effects on visual numerosity perception can also be produced by adapting to quantities in other sensory modalities (Anobile et al., 2021; Arrighi et al., 2014; Togoli & Arrighi, 2021), though these cross-modal adaptation effects are weaker than effects of adaptation to visual numerosity itself. Finally, beyond adaptation, numerosity estimation is reduced when individual items are connected by bars (He et al., 2009). This effect is not present in the earlier visual responses to numerosity (Fornaciai & Park, 2018a) and cannot be explained by changes in the spatial frequency domain contrast of the displays (Paul et al., 2022), so at least some effects on numerosity perception depend on later stages. We therefore propose that neural effects of at many stages of numerosity processing contribute to perceptual numerosity adaptation effects. Neural populations in many areas represent information about numerosity in either their monotonic or tuned responses, with hierarchical processing of each response across many stages (Harvey & Dumoulin, 2017; Paul et al., 2022) and tuned responses likely being derived from monotonic responses (Kim et al., 2021; Zorzi & Testolin, 2018). As adaptation may be best understood as a property of all neural responses, we can expect adaptation effects at all of these stages, with effects at one stage likely being inherited by the next.

Our results do not convincingly demonstrate that adaptation effects on early visual monotonic responses ultimately cause adaptation effects on numerosity-tuned responses. Indeed, it is not yet clear that early visual monotonic responses are required to produce numerosity-tuned response. Nevertheless, several findings suggest that adaptation effects on numerosity-tuned responses are inherited in part from effects on early visual contrast representations. First, almost all visual inputs to the cortex come through the primary visual cortex, which represents image features by contrast-driven responses in the spatial frequency domain. There is no other pathway through which numerosity tuned neurons could be activated by visual stimuli. Second, computational models for the derivation of numerosity-tuned responses (Dehaene & Changeux, 1993; Kim et al., 2021; Verguts & Fias, 2004; Zorzi & Testolin, 2018) generally rely on an intermediate stage with monotonic responses to numerosity. We have previously shown that the monotonic responses to numerosity shown by two very different neural network models (Kim et al., 2021; Stoianov & Zorzi, 2012) are better predicted by early visual responses to contrast (Paul et al., 2022). Changing the early visual contrast representation seems likely to change any response derived from this representation.

Adaptation often relies on presenting the same stimulus state repeatedly or over an extended period. However, our changing adaptor condition suppressed the early visual response to numerosity to an intermediate extent although it did not repeatedly present the same numerosity like the low and high adaptor conditions. This may be understood in as an interaction with a monotonic response to numerosity. If suppression of the early visual response depends on the level of recent early visual activity, the changing adaptor condition would be expected to produce an intermediate average level of activity, because the presented numerosity is always between the low and high adaptor. We would expect neural adaptation effects on numerosity-tuned responses to work quite differently because they would suppress the response similar throughout the response curve, not at a specific numerosity.

Indeed, all our conditions only presents the adaptor very briefly (and typically once) before each presentation of a changing numerosity, although many times over different presentations of changing numerosities. Is this sufficient to produce repulsive numerosity adaptation effects in perception? Or does this instead produce attractive serial dependence effects that occur when single presentations of a particular numerosity bias perception of the numerosity in the next presentation (Cicchini et al., 2014; Fornaciai & Park, 2018b)? We have previously shown that the stimulus timing used here produces a clear repulsive adaptation effects (Tsouli et al., 2021). Previous results also shown repulsive adaptation effects with brief adaptor presentations (Aagten-Murphy & Burr, 2016). Again here, these brief but frequent presentations, although separated by changing numerosities, would be expected to affect the average level of recent activity in the early visual cortex that we propose underlies the effects we observe.

Functionally, adaptation is usually proposed to adapt perception to the context of recently seen sensory stimuli, thereby increasing sensitivity in the stimulus range we are currently working with by increasing discriminability around the adapted range (Grzywacz & Balboa, 2002). Seeing contrast adaptation as a fundamental contributor to numerosity adaption instead suggests numerosity adaptation’s functional role may be to help to separate numerosity from contrast. Both numerosity and the contrast between items and their background (i.e. item contrast) similarly affect Fourier power in the spatial frequency domain (Paul et al., 2022). An image can have greater Fourier power because it contains more items or greater item contrast. To determine numerosity, we need to normalize the image contrast for item contrast. Indeed, responses in V1 are strongly contrast-dependent, while responses in the first areas showing numerosity-tuned responses (visual field maps TO1 and TO2, i.e. area hMT+) are minimally affected by item contrast (Kastner et al., 2004). Therefore, under normal circumstances, contrast adaptation may serve to normalize item contrast by considering the contrast of recently viewed items, and thereby yield a contrast-invariant representation to numerosity. However, during the unusual circumstances of numerosity adaptation, numerosity affects image contrast while item contrast is held constant. This may thereby disrupt this normalization process, leading to inaccurate numerosity perception. This view sees mechanisms of numerosity adaptation as inherent to the process of numerosity estimation itself, rather than an adaptive aspect of numerosity perception. These views are not mutually exclusive.

## Conclusions

The current results show a central role for early visual cortex in the neural basis of numerosity adaptation, increasing in strength through the visual processing hierarchy. These early visual effects may be inherited by later numerosity-tuned neural populations, with separate neural adaptation effects also likely acting in numerosity-tuned stages. Therefore, the neural basis of numerosity adaptation likely involves effects at all levels of numerosity processing. Together, these pervasive neural effects throughout the brain seem likely to underlie the strong and multi-faceted perceptual effects of numerosity adaptation.

## Acknowledgements

This work was supported by the Netherlands Organization for Scientific Research (452.17.012 to B.M.H.) and the Chinese Scholarship Council (202208510065 to L.Z.)

## Author contributions

Conceptualization, L.Z. , S.D. and B.M.H.; methodology, L.Z. and B.M.H.; software, L.Z. and B.M.H.; validation, L.Z., E.H. and B.M.H.; formal analysis, L.Z., E.H. and B.M.H.; investigation, L.Z. and B.M.H.; data curation, L.Z. and B.M.H.; writing – original draft, L.Z. and B.M.H.; writing – review & editing, L.Z., E.H., Y.W., S.G, S.D. and B.M.H.; visualization, L.Z., E.H. and B.M.H.; supervision, S.G., S.D. and B.M.H.; project administration, B.M.H.; funding acquisition, B.M.H.

## Method

### Participants

We recruited eight human participants (five male, three female; age range 26-52 years). One was left-handed. All were well educated, with good mathematical abilities, and had normal or corrected-to-normal visual acuity. All gave written informed consent. All experimental procedures were approved by the ethics committee of University Medical Center Utrecht (protocol number 09/350).

### Numerosity stimuli

We used MATLAB (MathWorks, Inc.) and the Psychophysics Toolbox (Brainard, 1997; Kleiner et al., 2007) to generate and present experimental stimuli similar to our past studies (Harvey et al., 2013, 2015; Harvey & Dumoulin, 2017; Tsouli et al., 2021). The numerosity stimuli were presented on a 69.84 × 39.29 cm LCD screen (Cambridge Research Systems) positioned behind the MRI bore. Participants were required to lie still and view the display through a mirror attached to the head coil. The total distance from the attached mirror to the display screen was 220 cm and the display resolution was 1920 × 1080 pixels.

Two large, thin and red cross lines were presented in the entire display to aid accurate fixation at the cross intersection in the center of the display. All items in the numerosity stimuli were positioned pseudo-randomly and limited within a circle centered on the fixation of 0.75° of visual angle (radius), minimizing the extent of the numerosity pattern, allowing it to be viewed without eye movements, and falling within the population receptive field of fMRI recording site responding to the central visual field. The pseudo-random positions of these items were constrained so that items were evenly spaced throughout this limited circle, avoiding perceptual grouping. Each numerosity stimulus presentation contained a new pseudo-random dot pattern. We kept the total surface area of all display items constant regardless of numerosity, so that display luminance was unaffected by numerosity.

In all conditions, the numerosities 1 through 7 and 20 were presented as dots on a gray background (Fig. 1). Each numerosity stimulus was presented briefly (300 ms) to ensure participants had no time to sequentially count the dots. The numerosity stimulus was followed by an interstimulus interval (ISI) of 400 ms showing a uniform gray background, then the next numerosity stimulus. In each 1400 ms (one fMRI volume acquisition, TR), we first showed an adaptor numerosity, which differed between three conditions, then a changing numerosity which was the same for all three conditions.

The changing numerosities varied from 1 through 7, with a baseline of 20 dots. For the changing numerosities 1-7, this repeated three times (across three TRs) before the numerosity changed, to ensure strong fMRI responses and allow enough time to distinguish the hemodynamic responses to different numerosities. When the changing numerosity 20 this was repeated 12 times (across 12 TRs) to better distinguish between numerosity-tuned and monotonic responses. A monotonically increasing response to numerosity should have a high amplitude during this period. However, a tuned response with a numerosity preference far below 20 (Cai et al., 2021) should have a lower amplitude during this period because a numerosity of 20 dots should be well outside of the range that elicits strong responses. This also allowed us to distinguish neural populations with very small tuning widths which never responded to the changing numerosities 1 through 7, and populations with very large tuning widths which always responded to these numerosities (Harvey et al., 2013).

In the low and high adaptor conditions, the alternating adaptor numerosity was held constant at 1 and 20 respectively. In the changing adaptor condition, the same numerosities were shown in the adaptor as the changing numerosity.

The changing numerosity stimuli were first presented in ascending order (1 to 7) for 4.2 s (3 TRs) each, next followed by 16.8 s (12 TRs) where the stimulus contained 20 dots, then followed by the numerosities in descending order (7 to 1) for 4.2 s (3 TRs) each, finally followed by another same long period of 20 dots. This sequence was repeated four times in each scanning run, resulting in a run duration of 369.6s. Therefore, each of the changing numerosity stimuli 1 through 7 was shown for a total of 24 times in each functional run. In the changing adaptor condition, these changing numerosities were also shown as the adaptor, adding another 24 times in each run.

The dots showing both the adaptor and changing numerosities were shown in black in 90% of dot presentations, while in the remaining 10%, the dots were shown in white (Fig. 1). Participants were instructed to press a button when the dots were shown in white instead of black (which is very easy at all numerosities) to ensure that they were paying attention to the stimuli during fMRI acquisition. No numerosity judgments were required.

### Visual field mapping stimuli

In a separate scanning session, visual field mapping was used to delineate visual field maps and determine the position selectivity of our recording sites, following protocols described previously (Dumoulin & Wandell, 2008; Harvey & Dumoulin, 2011; Paul et al., 2022). Briefly, a bar filled with a moving checkerboard pattern stepped across a 6.35° (radius) circle in the display center in eight (cardinal and diagonal) directions. Participants fixated the same central fixation cross, pressing a button when this changed color to ensure fixation and attention.

### MRI data collection

#### fMRI acquisition procedure

We acquired MRI data on a 7T Philips Achieva scanner for a previously published study (Tsouli et al., 2021). Similar acquisition protocols are described fully in other previous studies (Harvey et al., 2015; Harvey & Dumoulin, 2017). Briefly, we acquired T1-weighted anatomical scans, automatically segmented these with Freesurfer (http://freesurfer.net), then manually edited labels to minimize segmentation errors using ITK-SNAP (http://www.itksnap.org/). This provided a highly accurate cortical surface model at the grey-white matter border to the characterize cortical organization of the measured responses. Functional T2*-weighted 2D echo planar images were acquired using multiband acquisition (multiband factor: 2) and anterior-posterior encoding, and a 32-channel head coil, at a resolution of 1.77 × 1.77 × 1.75 mm, with a field of view of 227 × 227 × 70 mm. The TR was 1400 ms, echo time (TE) was 25 ms, and flip angle was 70°. Functional runs were each 273 time frames (382.2 s) in duration, of which the first 9 time frames (12.6 s) were discarded to ensure the signal was at steady state.

Three scanning sessions were required for each participant. In each scanning session, 3 functional runs were acquired for the changing adaptor condition (9 runs in total, total duration: 57 min 20 s) and 3-4 runs for the low and high adaptor conditions (in total 10 runs each for these adaptation conditions in total, total duration: 63 min 42 s; with the exception of one participant where 9 runs were acquired for each condition due to technical issues). The additional run we acquired for the low and high adaptor conditions helped ensure strong fMRI responses, because the changing numerosity stimuli were presented less frequently due to the interleaved adaptor stimuli. The order of the conditions was counterbalanced across runs within and between participants. Moreover, in each session we acquired a top-up scan recorded with the opposite phase-encoding direction to correct for image distortion in the gradient encoding direction (Andersson et al., 2003).

### fMRI preprocessing

The functional data was co-registered to the anatomical space using AFNI (afni.nimh.nih.gov; Cox, 1996) as described previously (Paul et al., 2022; Tsouli et al., 2021; van Ackooij et al., 2022). A single transformation matrix was constructed, incorporating all the steps from the raw data to the cortical surface model to reduce the number of interpolation steps to one. For the fMRI data, we first applied motion correction to the functional data (3dvolreg). We also applied motion correction to the images that were acquired using opposing phase-encoding direction, then determined the distortion transformation between these and the functional runs (3dQwarp) to correct for spatial distortions in the functional scans (3dNwarpApply). Then we determined the transformation that co-registers this functional data to the T1 with the same resolution, position and orientation as the functional data (3dvolreg). We finally determined the transformation from this T1 image to a higher resolution (1 mm isotropic) whole-brain T1 image (3dUnifize, 3dAllineate). We applied the product of all these transformations for every functional volume to transform our functional data to the whole-brain T1 anatomy. We repeated this for each fMRI session to transform all their data to the same anatomical space. We then imported these data into Vistasoft’s mrVista framework (github.com/vistalab/vistasoft) for analysis and model fitting. For each adaptation condition, the time series of separate scans were then averaged together, resulting in a very high signal-to-noise ratio.

#### fMRI Data Analysis

##### Neural response models for responses to numerosity

For each fMRI recording site (voxel) we interpret the fMRI responses to the numerosity stimuli using two neural response models: a numerosity-tuned population receptive field (pRF) model (Dumoulin & Wandell, 2008; Harvey et al., 2013, 2015; Harvey & Dumoulin, 2017) and a monotonic response model (DeWind et al., 2019; Park et al., 2015; Paul et al., 2022). These each describe the recording site’s response using a small set of parameters that we can then compare between adaptation conditions.

For the monotonic response model, the predicted neural response at each recording site is proportional to the logarithm of the aggregate Fourier power (in the spatial frequency domain) of the displays with each numerosity (Paul et al., 2022), shown at each time point. We convolved this neural response time course with an HRF to give an fMRI response time course prediction. In each adaptor condition, we used a general linear model to compare this prediction to the fMRI response time course at each recording site. This determined the slope of the relationship between the prediction and the response (proportional to the neural response amplitude, following a positive or negative relationship), together with the response variance explained by this scaled prediction. As we have previously shown (Paul et al., 2022), this contrast-driven response model is closely but nonlinearly related to a monotonic response to the logarithm of the presented numerosity in each display. However, it predicts the responses of the early visual cortex and neural network to numerosity displays more closely than numerosity does. We also repeated our analyses using a model describing a monotonic response to the logarithm of the presented numerosity at each time point, giving very similar results.

The numerosity-tuned pRF model describes the aggregate tuning of neural populations in each record site using a logarithmic Gaussian function with two free parameters: preferred numerosity (mean of the Gaussian function) and tuning width (standard deviation of the Gaussian in logarithmic numerosity space). We started by generating a large candidate set of combinations of preferred numerosity and tuning width. For every candidate combination, we predicted a neural response time course as the amplitude of the candidate neural response function at each time point’s presented numerosity. We then convolved this candidate neural response time course with a hemodynamic response function (HRF), giving a corresponding candidate fMRI response time course prediction. For each fMRI recording site and stimulus condition, we chose the fMRI response time course prediction that most closely followed the recorded response time series (by minimizing the sum of squared errors between the predicted and observed fMRI time series). We then took the parameter combination that generated this fMRI response time course prediction, together with the goodness of fit of this prediction, for further analysis. We quantify this goodness of fit as the variance explained by the model, i.e. R^2^, the proportion of the variance of the fMRI response time course that is outside the residual of the fit model.

In modelling the responses to the adaption conditions, we fit two different models. One model included only the aggregate Fourier power of the changing numerosity displays when making the predictions of fMRI responses, while the other included both the adaptor display and the changing numerosity display. We used the latter model for subsequent analyses as it was a complete description of the presented stimulus. However, the stimulus was designed so that both models would produce closely related predictions and parameter estimates. In the high and low adaptor conditions, the adaptor presented a constant numerosity throughout the run. This should produce a constant response throughout the run in both the monotonic and tuned response model. In such general linear modeling frameworks, this adaptor then adds a constant component to the predicted response. FMRI data has an arbitrary baseline that is anyway captured by another constant component in both models (which we do not analyze), so any further constant component contributes to that baseline without affecting other model parameters. In the changing adaptor condition, the adaptor presented the same numerosities as the changing numerosity that our models’ responses follow. In general linear models, this doubles the amplitude of the predicted response to the changing numerosity. This therefore halves the scaling between the fMRI response and the predicted response to the changing numerosity display, compared to a model that does not consider the response to the adaptor display. Considering this difference between adaptation conditions thereby makes the amplitudes of responses to the changing numerosity straightforwardly comparable between adaptation conditions. Any changes in fMRI responses between adaptation conditions can only arise through non-linear interactions between response to the adaptor and the changing numerosity stimuli.

It is important for further analyses to distinguish between monotonically increasing and numerosity-tuned responses. We used the changing adaptor condition to identify responses to changes in numerosity as we have used this stimulus design in previous studies (Harvey et al., 2013; Harvey & Dumoulin, 2017; Paul et al., 2022; Tsouli et al., 2021) and it maximizes neural response amplitude and the goodness of fit of our models. We fit both monotonic and tuned models to the averages of the odd and even numbered scans in this condition. We then evaluated the response predictions of the both resulting models on the complementary half of the data (i.e. cross-validation) because the tuned model is fit from a larger set of predictions which follow more complex functions. During this evaluation, we allowed the response predictions to rescale in amplitude (but not change sign) between fitting and evaluation because the complementary halves of the data were often acquired in different scanning sessions, which can arbitrarily differ in fMRI signal amplitude. We then computed the residual sum of squared errors between the responses and predictions across both halves and for each voxel chose the model with the lower residual.

A numerosity-tuned response can be clearly identified when the preferred numerosity is within the range of the changing numerosities 1 through 7, because this shows the response amplitude decrease for higher numerosities. Therefore, our numerosity-tuned pRF models make and test predictions outside of this range to show that preferred numerosity estimates within this range predict responses better than functions with a preferred numerosity outside of this range. A monotonic response can be clearly identified when a monotonic response model fits better than a numerosity-tuned model. However, voxels that fit slightly better by a numerosity-tuned model with a numerosity preference above 7 are also likely to reflect monotonic responses, because our previous experiments using a larger numerosity range demonstrate that very few voxels show numerosity-tuned responses with preferences above 7 (Cai et al., 2021). We therefore also use monotonic models of voxels where the numerosity-tuned model estimates a numerosity preference above 7.

Moreover, we also exclude from further analysis of numerosity-tuned pRF models the recording sites for which the response models in the changing adaptor condition explained less than 0.2 of response variance.

##### Neural response models for visual field position and definition of visual field maps

We localized monotonic responses to the area around the occipital pole, the location of the visual field maps of the early visual cortex (DeWind et al., 2019; Park et al., 2015; Paul et al., 2022). We therefore asked how adaptation effects on monotonic responses are localized in these early visual field maps. We fit the responses to the visual field mapping stimuli using a standard visual spatial pRF analysis (Dumoulin & Wandell, 2008; Harvey & Dumoulin, 2011). We defined visual field maps borders based on the reversals in the cortical progression of the polar angle of voxels’ visual field position preferences, manually identifying these on an inflated rendering of each participant’s cortical surface (Dumoulin & Wandell, 2008; Harvey & Dumoulin, 2011). These formed our main regions of interest. As well as the early visual field maps (V1, V2, V3, hV4), we also identified mid-level visual field maps (LO1, LO2 and V3A/B) which showed monotonically-responding recording sites in some hemispheres.

### Comparisons and statistics

In order to quantify the change in monotonic response amplitudes between different adaptation conditions, we analyzed the parameters of monotonic models fit to the responses of recording sites in each the early visual field maps (V1-V3, hV4, V3A/B, LO1, LO2). Specifically, we compared the slope of the relationship between the monotonic response prediction and the recorded response, i.e. the increase in amplitude of the neural response underlying the fMRI signal when the aggregate Fourier power of the changing numerosity display increases by one (Fig. 3B). We also repeated this using a log(numerosity) response model, where the slope parameter reflects the increase in amplitude of the neural response underlying the fMRI signal when the logarithm of the presented numerosity increases by one. This gave very similar results.

To make these comparisons between monotonic responses in the different adaptation conditions, we first take all the recording sites within a visual field maps and extract their preferred visual field positions from the visual field position response models. For each recording site, we then extracted the fit slope from the monotonic numerosity response models for each adaptation. Within each visual field maps, we then select recording sites that meet the following criteria for further analysis: (1) where the preferred visual field position’s eccentricity is below 1°, i.e, recording sites whose visual spatial population receptive field include the numerosity stimulus area; (2) the slope of the monotonic model in the control condition is positive, so response amplitudes increase with numerosity; and (3) the model variance explained in the control condition is at least 0.1, corresponding to under 5% probability of observing these responses by chance. We then calculated the average slope among the selected voxels in each visual field map in each hemisphere (i.e., in each visual field map example) for each adaptation condition.

In subsequent analyses, for each visual field map, we use the resulting slope in each visual field map example as independent measures. We first tested for significant differences between these slopes and variance explained using the Wilcoxon signed rank test, where the values for each hemisphere in one adaptation condition and paired with the values from the same visual field map example in the other adaptation conditions, i.e., we tested whether the difference in these visual field map examples’ slopes between these two adaptation conditions was significantly above zero. As we performed this comparison separately for each visual field map, we performed a false discovery rate (FDR) correction (Benjamini & Hochberg, 1995) on the resulting probability estimates, taking all visual field maps into account.

We also ask whether the strength of the adaptation effect on the monotonic model slope differed between visual field maps. This is complicated by the fact that, within each adaptation condition, the slopes shows clear differences between visual field maps, making it difficult to interpret any changes between adaptation conditions. We would expect a visual field map with a high slope or high variance explained to be able to decrease this slope more (in absolute values) with adaptation.

We therefore calculated the change in slope between the low and high adaptor conditions, between the low adaptor and changing adaptor conditions, and between the changing adaptor and high adaptor conditions. In each case, we divided this decrease in slope by the slope in the changing adaptor condition to give a proportion by which the slope changed that was comparable between pairs of conditions. Having calculated the proportion by which the slope decreased from these three comparisons in each visual field map example, we performed separate two-factor ANOVAs for each pair of conditions (factors: visual field map and participant) to test whether the proportional decrease in slope differs between visual field maps. These are corrected for multiple comparisons by using Tukey’s honestly significant difference test (Tukey, 1949), which gives the marginal means and confidence intervals shown in Fig. 4F.

